# The MLL1–MENIN complex preserves CD8 T cell memory through a TOX–BTLA-TCF1 axis

**DOI:** 10.64898/2026.04.03.715913

**Authors:** Bo Chin Chiu

**Affiliations:** From the Department of Pathology, University of Michigan Medical School, Ann Arbor, Michigan

## Abstract

Immunological memory depends on the maintenance of stem cell-like memory CD8 T cells, which require sustained expression of the transcription factors TCF1. Here, I identify MLL1 as a key regulator of CD8 T cell memory. In activated T cells, MLL1 sustains *Tox* transcription through interaction with MENIN, thereby maintaining BTLA expression and restraining cytokine-driven AKT activation. Loss of MLL1 or disruption of the MLL1–MENIN interaction accelerates AKT-driven loss of TCF1, leading to impaired memory potential. MLL1-deficient T cells fail to reconstitute lymphopenic hosts and are unable to mediate graft-versus-host disease, while exhibiting increased expansion of virtual memory T cells. Unexpectedly, MLL1 regulates *Tox*, *Btla* and *Tcf7* independently of its methyltransferase activity and MOF-mediated H4K16 acetylation. These findings define a pathway in which the MLL1–MENIN complex restrains cytokine signaling to preserve CD8 T cell memory and identify a noncanonical function of MLL1 in transcriptional maintenance.

## Introduction

In response to infection, antigen-specific T cells undergo asymmetric division to generate short-lived effector cells that provide immediate protection and long-lived memory cells that confer durable immunity against reinfection ^1–6^. Long-lived memory T cells exhibit regenerative capacity similar to that of hematopoietic stem cells and are therefore often referred to as stem cell–like memory T cells ^7,8^. The development of these cells depends on the transcription factor TCF1 (encoded by Tcf7). In the absence of TCF1, activated T cells preferentially differentiate into short-lived effector cells ^9–11^.

Naïve and long-lived memory T cells express high levels of TCF1, but its expression decreases following antigen stimulation. This reduction is regulated by T cell receptor (TCR) and cytokine signaling, with stronger stimulation resulting in lower TCF1 levels ^9,10,12,13^. This mechanism enables effector differentiation to be tailored to the level as well as the nature of the antigen. However, regulatory pathways must exist to restrain effector differentiation and prevent the complete loss of the memory T cell pool. One such mechanism involves TOX, which limits TCR and cytokine signaling by upregulating co-inhibitory receptors such as PD-1 and BTLA ^14–17^. In the absence of TOX, activated T cells undergo accelerated differentiation and fail to persist. However, the mechanisms that maintain *Tox* expression during T cell activation remain poorly understood.

The CD8⁺ T cell compartment can be broadly divided into naïve and memory populations based on CD44 expression, with naïve and memory cells expressing low and high levels, respectively ^18^. The memory population can be further subdivided into true memory (TM) and virtual memory (VM) T cells. TM cells arise from naïve cells following antigen stimulation, whereas VM cells are antigen-inexperienced and develop in response to cytokine signaling ^19,20^. In laboratory mice with limited antigen exposure, the majority of memory CD8⁺ T cells are VM cells ^21^. Thymic involution begins early in life and accelerates after puberty ^22^, leading to reduced output of naïve T cells. Nevertheless, the total number of peripheral T cells remains relatively constant due to the progressive expansion of memory T cells ^23,24^. In mice, cytokine-driven expansion of VM cells accounts for most of the age-associated memory T cell expansion^21^.

MLL1 is a ubiquitously expressed DNA-binding protein that co-localizes with RNA polymerase II at the promoters of actively transcribed genes ^25,26^. MLL1 has been shown to regulate gene expression through multiple mechanisms, including catalyzing H3K4 trimethylation at promoters of actively transcribed genes and recruiting MOF to mediate H4K16 acetylation independently of its methyltransferase activity ^27–30^. Additionally, MLL1 has been implicated in mitotic bookmarking ^30^. Whether such mechanisms operate in T cells, and how they might influence T cell fate decisions, remains unclear. In this study, using T cell–specific MLL1-deficient mice, I show that MLL1 preserves T cell memory by maintaining *Tox* transcription through interaction with MENIN. I demonstrate that MLL1 restrains cytokine-driven differentiation by sustaining a TOX–BTLA pathway, thereby preserving TCF1 expression and memory potential. Consistent with the phenotype of BTLA-deficient T cells ^31^, MLL1-deficient T cells fail to reconstitute T cell–deficient mice and are unable to induce graft-versus-host disease. Furthermore, MLL1-deficient VM T cells undergo accelerated expansion, mirroring their BTLA-deficient counterparts. Unexpectedly, these functions of MLL1 occurs independently of its canonical methyltransferase activity and H4K16 acetylation, supporting a non-catalytic role in stabilizing transcriptional programs during T cell activation.

## Results

### MLL1 is required for T cell regenerative capacity

Long-lived memory T cells possess stem cell-like regenerative capacity, enabling reconstitution of antigen-specific immunity ^7,8^. Given the essential role of MLL1 in hematopoietic stem cell maintenance ^32,33^, I asked whether MLL1 similarly regulates T cell regenerative potential. To address this, I used T cell-specific Mll1-deficient (Mll1KO) mice that were generated by deleting exons 7 and 8 in double-positive thymocytes using CD4-Cre. This strategy results in complete loss of MLL1 protein without affecting T cell development or peripheral homeostasis ^33^. I then assessed the regenerative capacity of Mll1KO CD8⁺ T cells using a competitive adoptive transfer model. Congenically marked Mll1KO (CD45.2⁺) and wild-type (WT; CD45.1⁺) CD8⁺ T cells were mixed at a 1:1 ratio, activated in vitro, and transferred into lymphopenic Rag1-deficient recipients (Figure 1A). Two weeks after transfer, donor-derived T cells were analyzed across multiple tissues. Strikingly, the vast majority of donor T cells in recipient mice were derived from WT cells, with minimal contribution from Mll1KO cells across all organs examined, which included spleen, blood, peripheral lymph nodes (PLN) and mesenteric lymph nodes (MLN) (Figure B and C). These findings demonstrate that MLL1 is required for the regenerative capacity of activated T cells, analogous to its role in hematopoietic stem cells.

**Figure 1.**
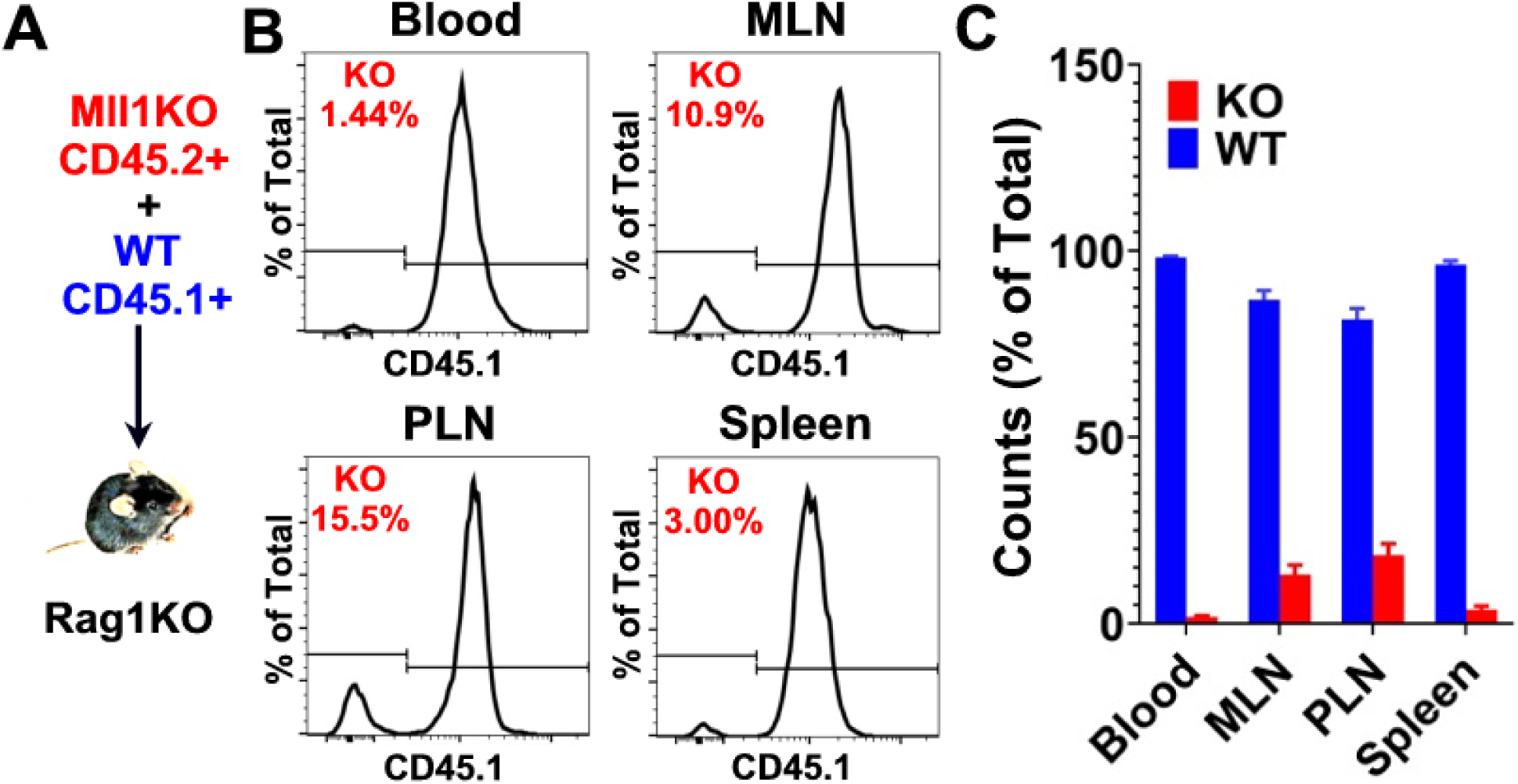
MLL1 is required for T cell regenerative capacity. Congenically marked CD8⁺ T cells from Mll1KO (CD45.2⁺) and WT (CD45.1⁺) mice were mixed at 1:1 ratio, activated in vitro for 4 days and used to reconstitute T cells deficient (Rag1KO) mice. Donor-derived cells in recipient mice were analyzed by flow cytometry 2 weeks after transfer. (A) Schematic illustration of the reconstitution experiment. (B, C) Peripheral blood, mesenteric lymph nodes (MLN), peripheral lymph nodes (PLN), and spleen were collected and analyzed by flow cytometry. (B) Representative flow cytometry plots of gated CD8⁺ T cells, showing the frequency of Mll1KO cells, identified by the absence of CD45.1 expression. (C) Summary of the percentages of Mll1KO and WT CD8⁺ T cells in the indicated tissues. Data are representative of five independent experiments.

### MLL1 maintains TCF1 and CD62L through interaction with MENIN

TCF1 and CD62L are co-expressed in T cells and mark populations with high regenerative potential ^9–11^. Loss of TCF1 and CD62L has been associated with diminished regenerative potential ^7,8^. To determine whether MLL1 regulates this program, I activated Mll1KO and WT CD8 T cells and assessed co-expression of TCF1 and CD62L. As shown in Figure 2A, approximately 50% of WT T cells co-expressed CD62L and TCF1 four days after activation. In contrast, a significantly lower proportion (∼27%) of Mll1KO T cells retained co-expression of these markers (Figure 2B).

**Figure 2.**
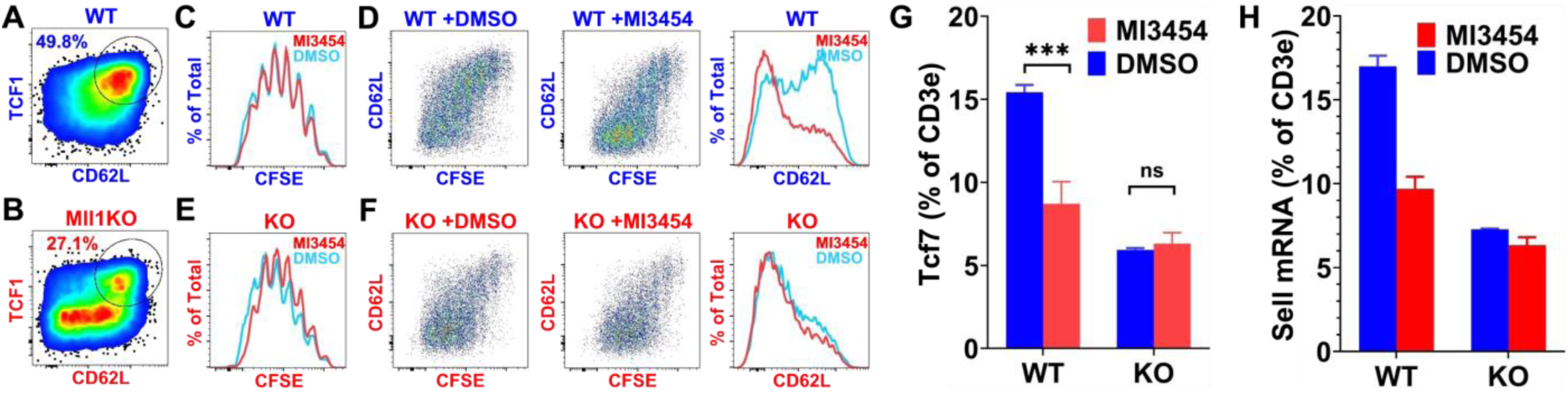
MLL1 maintains TCF1 and CD62L through interaction with MENIN. (A, B) CD8 T cells from WT (A) mice and their Mll1KO (B) littermates were activated in vitro with anti-CD3, anti-CD28 and IL-2. After 4 days, cells were collected and analyzed for TCF1 and CD62L expression by flow cytometry. Data are representative of three independent experiments. (C-F) CD8 T cells from WT (C, D) mice and their Mll1KO (E, F) littermates were labeled with CFSE and activated in the presence of MI3454 or DMSO. After 4 days, cells were analyzed for proliferation based on CFSE dilution (C, E) and for CD62L expression (D, F) by flow cytometry. Data are representative of two independent experiments. (G, H) CD8 T cells from WT mice and their Mll1KO littermates were activated in the presence of MI3454 or DMSO. After 4 days, cells were collected and analyzed for *Tcf7* (G) and *Sell* (H) transcript levels by RT-PCR. Data are representative of two independent experiments.

MLL1 is known to regulate gene transcription through interaction with MENIN. Consistent with this, MENIN-deficient T cells exhibit accelerated loss of CD62L following activation ^34–36^. To assess the role of the MLL1–MENIN interaction, I activated CD8⁺ T cells in vitro in the presence of MI-3454, an inhibitor that disrupts MENIN interactions, or DMSO as a control ^37^. Because CD62L loss is closely linked to cell division, cells were labeled with CFSE prior to activation to track proliferation. At concentrations ≥0.5 μM, MI-3454 inhibited cell division; therefore, 0.25 μM was used for subsequent experiments, as it had minimal impact on proliferation (Figure 2C). As expected, CD62L expression decreased progressively with each cell division (Figure 2D). Notably, MI-3454 treatment accelerated CD62L loss compared to control conditions, resulting in a higher proportion of CD62L⁻ cells. Although MI3454 was developed to block the interaction between MENIN and MLL1, it also blocks the interaction between MENIN and other MENIN-interacting partners, including MLL2 and JUND ^37^. To test whether its effect on CD62L was MLL1-dependent, Mll1KO T cells T cells were activated in the presence of MI-3454 or DMSO. MI-3454 did not affect proliferation in Mll1KO T cells (Figure 2E). Notably, the effect of MI-3454 on CD62L was abolished in Mll1KO cells (Figure 2F), indicating that it is MLL1-dependent. These findings indicate that MLL1 and MENIN cooperatively maintain CD62L expression in activated T cells.

Naïve T cells express high levels of TCF1, which decline following antigen stimulation. Loss of TCF1 results in differentiation into short-lived effector cells ^9,10^. Recent studies have shown that Mll1KO follicular helper T cells express reduced levels of TCF1 compared to WT cells ^38^, although the underlying mechanism remained unclear. Similarly, MENIN-deficient T cells also exhibit reduced TCF1 expression^34^. To determine whether MLL1 regulates TCF1 through MENIN, I activated WT and Mll1KO T cells in the presence of MI-3454 or DMSO and performed RT-PCR analysis. Consistent with the flow cytometry data (Figure 2A, B), activated Mll1KO T cells expressed lower levels of *Tcf7* compared to WT cells (Figure 2G). MI-3454 treatment significantly reduced *Tcf7* expression in WT T cells but had minimal effect in Mll1KO T cells. Similarly, expression of *Sell* (encoding CD62L) was reduced in Mll1KO T cells and decreased upon MI-3454 treatment in WT but not Mll1KO cells (Figure 2H). Together, these data demonstrate that MLL1 and MENIN function as a complex to maintain expression of genes associated with T cell memory.

### MLL1 regulates Tcf7 independently of its methyltransferase activity

MLL1 is a histone methyltransferase best known for catalyzing trimethylation of histone H3 lysine 4 (H3K4me3) ^27,39^. This activity depends on its interaction with WDR5, a scaffold protein that binds histone H3 and presents the H3K4 residue for methylation ^40–42^. The WIN site of WDR5 mediates interaction with arginine-containing motifs in multiple nuclear proteins, including MLL1. Pharmacological blockade of the WIN site displaces WDR5 from chromatin and can reduce gene transcription ^43^. To assess the role of MLL1 methyltransferase activity in regulating *Sell* and *Tcf7*, I treated activated wild-type (WT) T cells with MM-401, a peptidomimetic inhibitor of the WDR5 WIN site ^40,44^. MM-401 was used at 25 μM, as higher concentrations impaired cell division. At this concentration, MM-401 had minimal effects on proliferation (Figure 3A) and did not significantly alter *Sell* or *Tcf7* transcription (Figure 3C, D). Next, I used WDR5-IN-4, a more potent WIN site inhibitor ^45,46^. While concentrations ≥5 μM inhibited cell division, treatment at 2.5 μM had minimal impact on proliferation (Figure 3B). Unexpectedly, WDR5-IN-4 treatment increased, rather than decreased, *Sell* and *Tcf7* transcription (Figure 3C, D). These findings do not support a model in which MLL1 maintains *Sell* and *Tcf7* expression through H3K4me3 deposition at their gene loci.

**Figure 3.**
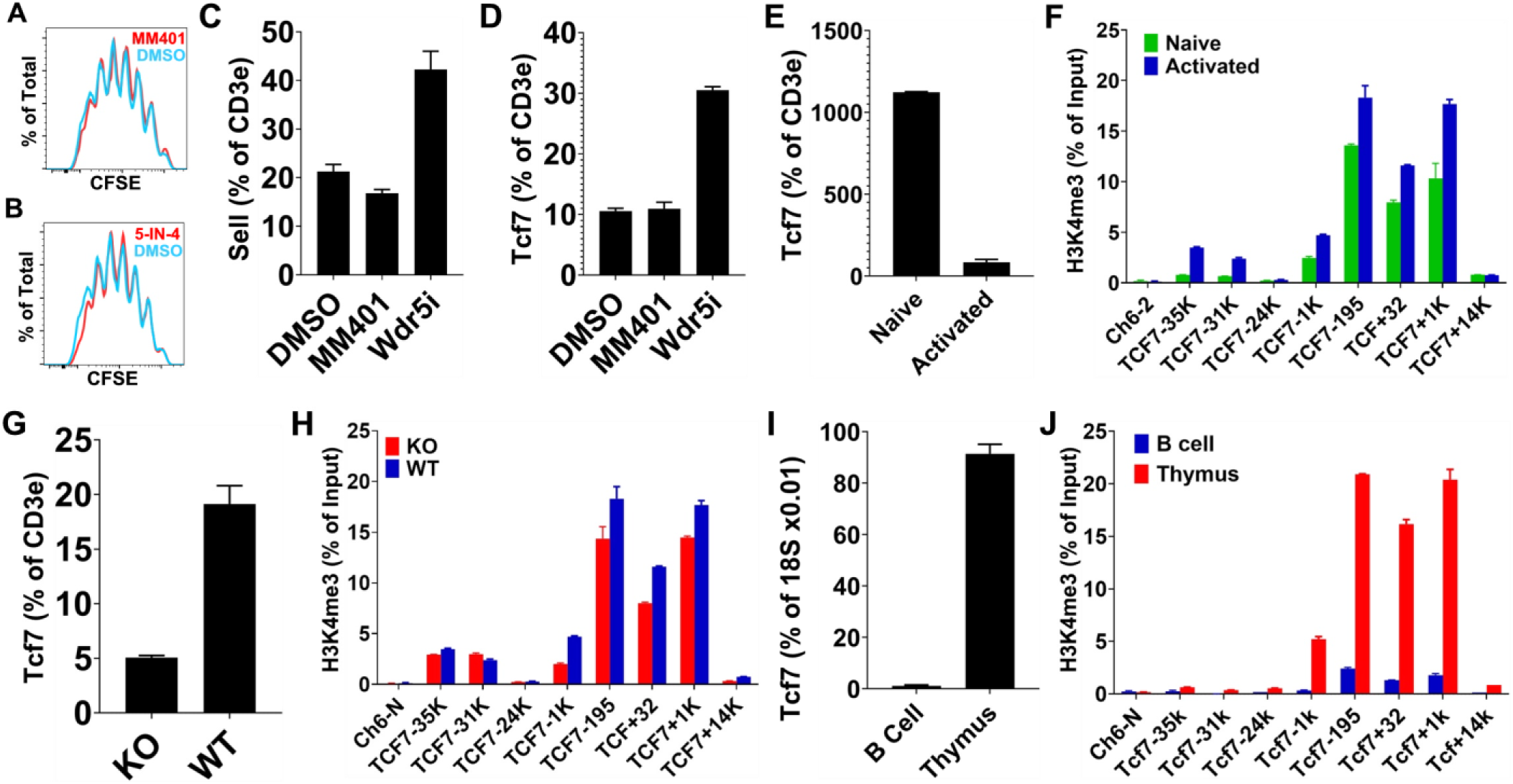
MLL1 regulates Tcf7 independently of its methyltransferase activity. (A–D) Wild-type (WT) T cells were labeled with CFSE and activated in the presence of MM-401, WDR5-IN-4, or DMSO. After 4 days, cells were analyzed by flow cytometry to assess the effects of MM-401 (A) and WDR5-IN-4 (B) on cell division (CFSE dilution), and by RT-PCR to determine the effects of these inhibitors on *Sell* (C) and *Tcf7* (D) transcription. Data are representative of two independent experiments. (E, F) Naïve CD8⁺ T cells were isolated from WT mice and activated in vitro for 4 days to generate activated T cells. Naïve and activated T cells were compared for *Tcf7* expression by RT-PCR (E) and for H3K4me3 enrichment at the *Tcf7* locus by ChIP-PCR (F). Data are representative of two independent experiments. (G, H) Mll1KO and WT T cells were activated in vitro and analyzed after 4 days for *Tcf7* transcription by RT-PCR (G) and H3K4me3 enrichment at the *Tcf7* locus by ChIP-PCR (H). Data are representative of three independent experiments. (I, J) Thymocytes and B cells were isolated from WT mice and compared for *Tcf7* transcription by RT-PCR (I) and H3K4me3 enrichment at the *Tcf7* locus by ChIP-PCR (J). Data are representative of two independent experiments.

Consistent with prior observations, *Tcf7* expression was substantially higher in naïve T cells than in activated T cells (Figure 3E). To examine the relationship between transcription and H3K4me3, I performed ChIP-PCR analysis of the *Tcf7* locus, including the transcription start site and previously identified enhancer regions (−35 kb, −31 kb, −24 kb, and +14 kb) ^47^. A gene desert region on chromosome 6 (Ch6-N) was used as a negative control. Surprisingly, H3K4me3 was highly enriched near the *Tcf7* transcription start site (−1 kb, −195 bp, +32 bp, and +1 kb) in both naïve and activated T cells (Figure 3F), despite marked differences in transcriptional output. These results suggest that *Tcf7* downregulation occurs independently of changes in H3K4me3 at this locus. Next, I compared activated Mll1KO and WT T cells. As expected, Mll1KO T cells expressed significantly lower levels of *Tcf7* (Figure 3G). However, H3K4me3 levels at the *Tcf7* locus were comparable between Mll1KO and WT cells (Figure 3H), indicating that MLL1 is not required for maintaining H3K4me3 at this site in activated T cells.

To further examine the relationship between *Tcf7* expression and H3K4me3, I compared thymocytes and B cells. *Tcf7* expression was more than 100-fold higher in thymocytes than in B cells, consistent with its role in T cell development ^10^. Although H3K4me3 enrichment at the *Tcf7* locus was substantially higher in thymocytes, detectable enrichment remained in B cells relative to the negative control (Figure 3J). Tcf7 is upregulated in hematopoietic stem cells and then downregulated upon differentiation ^48,49^. The fact that H3K4me3 is still enriched at the *Tcf7* locus in B cells suggests that loss of H3K4me3 is a slower process. These findings suggest that H3K4me3 does not instruct *Tcf7* transcriptional activity. Instead, H3K4me3 may reflect prior transcriptional activity, as proposed in recent studies ^50^. Together, these findings indicate that MLL1 maintains *Tcf7* transcription independently of its canonical methyltransferase activity in activated T cells.

### MLL1 maintains TCF1 by restraining cytokine-driven AKT signaling

In the absence of MENIN, mTORC1 and AKT signaling pathways are hyperactivated in activated T cells ^34–36^. Because *Tcf7* is regulated by both AKT and mTORC1 ^51,52^, I investigated the contribution of these pathways to MLL1-dependent regulation. T cells were activated in the presence of an AKT inhibitor (AKT1/2i) or rapamycin, an mTORC1 inhibitor. As shown in Figure 4A, inhibition of mTORC1 did not prevent MI-3454–mediated downregulation of *Tcf7*. In contrast, inhibition of AKT abrogated this effect, indicating that the MLL1–MENIN complex maintains *Tcf7* expression in an AKT-dependent but mTORC1-independent manner. Upon activation, AKT phosphorylates FOXO1 at conserved serine and threonine residues, promoting its nuclear export and degradation ^53^. FOXO1 is essential for maintaining T cell longevity; in its absence, activated T cells differentiate into short-lived effector cells ^53–56^. Notably, FOXO1 directly promotes transcription of genes associated with memory T cell fate, including *Tcf7*, *Sell* (CD62L), *Ccr7*, and *Il7r* ^51,53,57–61^. IL-7Rα expression enables responsiveness to IL-7, a key survival factor for T cells, while CD62L and CCR7 facilitate trafficking to lymphoid tissues where IL-7 is available ^62–65^. Together, these observations suggest that the MLL1–MENIN complex preserves memory T cell potential by restraining AKT activity and sustaining FOXO1 function.

**Figure 4.**
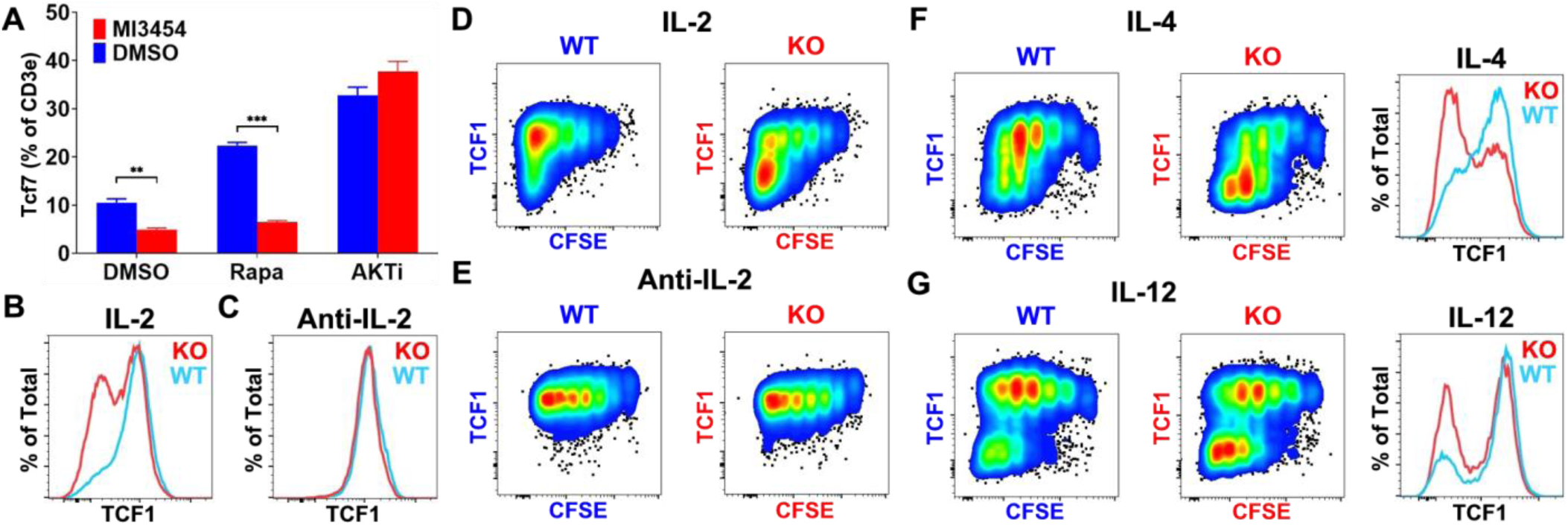
MLL1 maintains TCF1 by restraining cytokine-driven AKT signaling. (A) Wild-type (WT) T cells were activated in the presence of MI-3454, rapamycin, AKT1/2 inhibitor (AKTi), and/or DMSO. After 4 days, cells were collected and analyzed for *Tcf7* expression by RT-PCR. Data are representative of two independent experiments. (B, C) Mll1KO and WT T cells were activated in the presence of IL-2 (B) or neutralizing anti–IL-2 antibodies (C). After 4 days, cells were analyzed for TCF1 expression by flow cytometry. Data are representative of three independent experiments. (D–G) Mll1KO and WT T cells were labeled with CFSE and activated in the presence of IL-2 (D), anti–IL-2 (E), IL-4 (F), or IL-12 (G). After 4 days, cells were analyzed for TCF1 expression by flow cytometry. Data are representative of three independent experiments.

AKT activity is regulated by PDK1 and mTORC2. T cell receptor (TCR) and co-stimulatory signaling induce rapid but transient phosphorylation of AKT at T308 and S473, whereas sustained activation requires cytokine signaling ^53,66,67^. This mechanism enables T cells to balance effector and memory differentiation in response to both antigen load and cytokine milieu. IL-2 is a key regulator of this process, in part by sustaining AKT signaling ^6,68^. During T cell activation, the IL-2 receptor is asymmetrically partitioned, such that daughter cells inheriting higher receptor levels receive stronger IL-2 signals and preferentially differentiate into short-lived effector cells ^6,68^. To assess the role of cytokine signaling, I activated Mll1KO and WT T cells in the presence of IL-2 or neutralizing anti–IL-2 antibodies. Because IL-2 is required for T cell survival and expansion, IL-15 was included to support cell viability. As expected, in the presence of IL-2, a significantly greater proportion of Mll1KO T cells lost TCF1 expression compared to WT cells (Figure 4B). In contrast, under IL-2–neutralizing conditions, Mll1KO and WT T cells maintained similar levels of TCF1 (Figure 4C). These findings indicate that MLL1 preserves TCF1 expression by limiting IL-2–driven signaling.

To determine whether these effects were linked to proliferation, T cells were labeled with CFSE prior to activation. Mll1KO and WT T cells exhibited comparable proliferation under both IL-2–supplemented (Figure 4D) and IL-2–neutralized (Figure 4E) conditions. Notably, even in the presence of IL-2, both genotypes maintained TCF1 expression for at least three cell divisions, suggesting that sustained, rather than transient, AKT signaling is required for TCF1 downregulation. In addition to IL-2, AKT is activated by other cytokines, including IL-4 ^69^ and IL-12 ^70^. To determine whether MLL1 broadly restrains cytokine responsiveness, I activated Mll1KO and WT T cells in the presence of IL-4 or IL-12. Under both conditions, WT T cells maintained higher levels of TCF1 compared to Mll1KO cells (Figure 4F, G). Collectively, these data demonstrate that MLL1 maintains *Tcf7* expression by restraining cytokine-driven AKT signaling, independently of mTORC1.

### MLL1 maintains TOX in activated T cells

To identify upstream regulators of cytokine responsiveness, I performed RNA-seq analysis on CD8⁺ T cells activated in the presence of MI-3454 or DMSO. Compared to control cells, 457 genes were upregulated and 79 genes were downregulated by more than 1.5-fold in MI-3454–treated cells. The top 10 upregulated and downregulated genes are listed in Table 1. Notably, *Tox*, a key regulator that restrains effector T cell differentiation ^15,71^, was the most downregulated gene following MI-3454 treatment. Consistent with this, *Gzma* and *Cd244*, which are negatively regulated by TOX ^72^, were among the most upregulated genes, whereas *Sell*, *Il6ra*, and *Btla*, known TOX targets ^72^, were among the most downregulated. These data suggest that loss of *Tox* expression accounts for a substantial portion of the transcriptional changes induced by MI-3454.

**Table 1.**
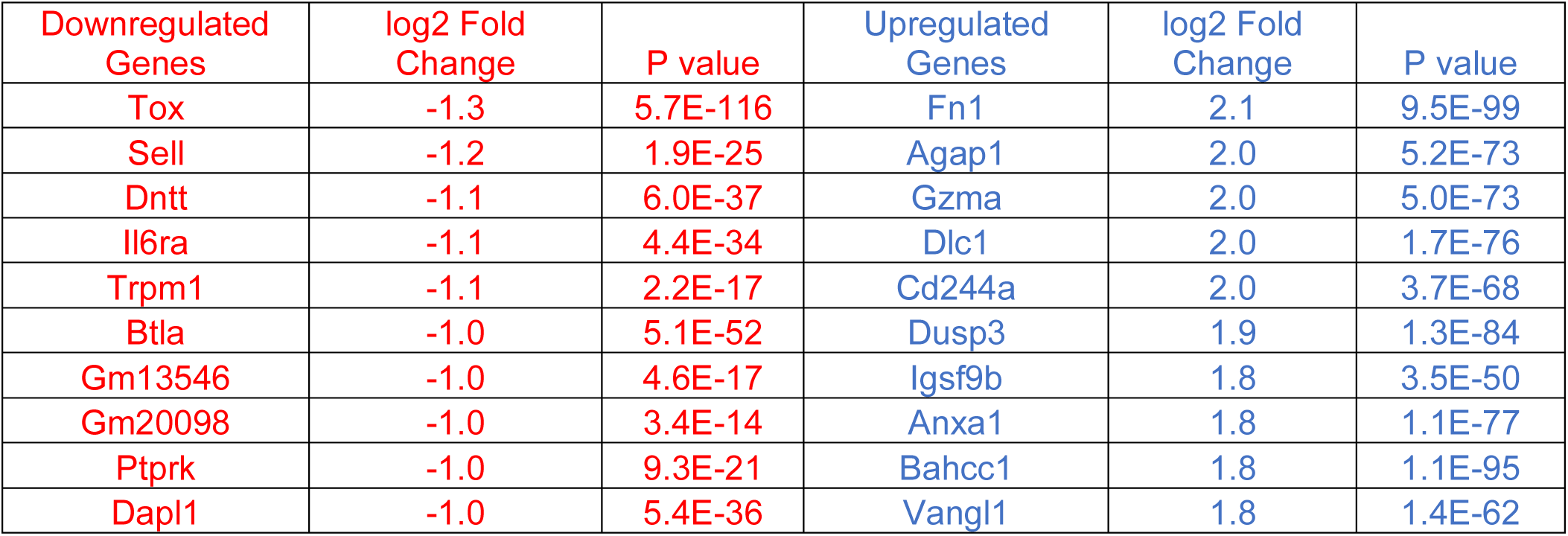
T cells from WT mice were activated in the presence of MI3454 or DMSO. After 4 days, the cells were collected and analyzed by RNA-seq.

To determine whether cytokine signaling contributes to *Tox* regulation, I activated WT T cells under IL-2–neutralizing conditions (anti–IL-2 + IL-15) in the presence of MI-3454 or DMSO. As shown in Figure 5A, MI-3454 treatment reduced *Tox* and *Btla* expression even in the absence of IL-2 signaling. Next, I activated T cells from Mll1KO mice and their WT littermates under IL-2–neutralizing conditions. Mll1KO T cells expressed lower levels of *Tox* and *Btla* than WT cells under these conditions (Figure 5B). These findings indicate that TOX and BTLA act upstream of cytokine–induced AKT activation, whereas TCF1 functions downstream.

**Figure 5.**
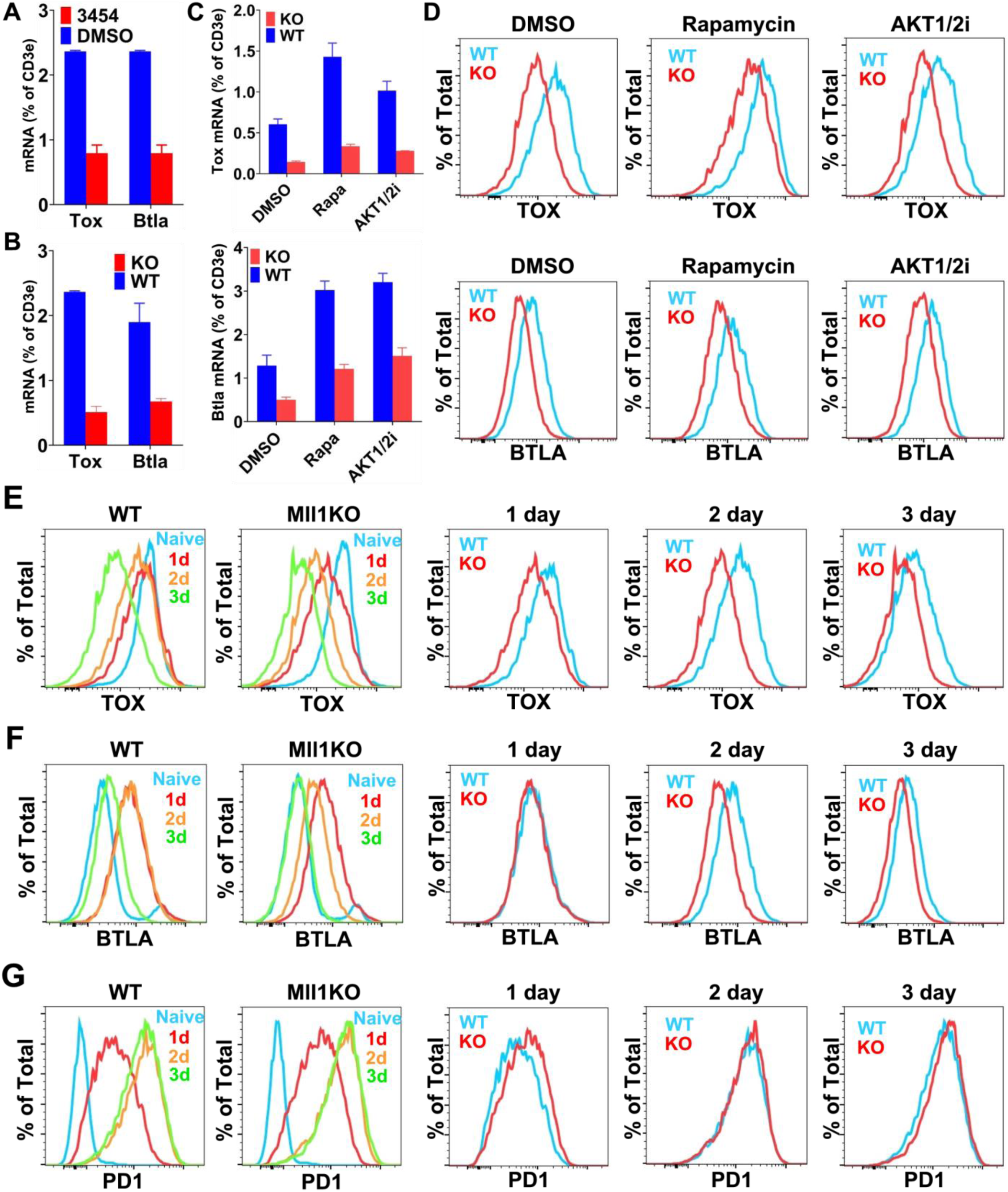
MLL1 maintains TOX in activated T cells. (A) Wild-type (WT) T cells were activated under IL-2–neutralizing conditions (anti–IL-2 + IL-15) in the presence of MI-3454 or DMSO. After 4 days, cells were collected and analyzed for *Tox* and *Btla* expression by RT-PCR. Data are representative of three independent experiments. (B) T cells from Mll1KO mice and their WT littermates were activated under IL-2–neutralizing conditions (anti–IL-2 + IL-15) in the presence of MI-3454 or DMSO. After 4 days, cells were collected and analyzed for *Tox* expression by RT-PCR. Data are representative of three independent experiments. (C, D) T cells from Mll1KO mice and their WT littermates were activated in vitro in the presence of rapamycin, AKT1/2i, or DMSO. After 4 days, cells were analyzed for *Tox* expression by RT-PCR (C) and for TOX protein levels by flow cytometry (D). Data are representative of two independent experiments. (E–G) T cells from Mll1KO mice and their WT littermates were activated in vitro. Cells were collected before activation (naïve) and at 1, 2, and 3 days post-activation, and analyzed by flow cytometry for TOX (E), BTLA (F), and PD-1 (G) expression. Data are representative of two independent experiments.

The *Tox* gene was originally identified in thymocytes, where its expression is dynamically regulated during development ^73^. Pre-selection double-positive thymocytes express low levels of *Tox*, which is rapidly upregulated upon positive selection and subsequently downregulated during maturation. In contrast, mature T cells downregulate *Tox* following TCR stimulation ^71^. At the transcriptional level, *Tox* is regulated by T-bet and EOMES, with T-bet represses, whereas EOMES promotes *Tox* expression ^71^. Both factors are regulated by mTORC1 signaling ^74^, such that increased mTORC1 activity is expected to reduce *Tox* transcription. Conversely, TOX has been shown to inhibit mTORC1 activity ^15,71^, suggesting a bidirectional regulatory relationship. To understand the relationship between TOX, mTORC1 and AKT, I activated WT and Mll1KO T cells in the presence of rapamycin or AKT1/2i. RT-PCR analysis showed that inhibition of either pathway increased *Tox* and *Btla* expression (Figure 5C). However, neither treatment rescued the reduced expression of these genes in Mll1KO T cells. Flow cytometry confirmed that TOX and BTLA protein levels remained lower in Mll1KO T cells compared to WT controls, even following inhibition of AKT or mTORC1 (Figure 5D). These results indicate that the heightened cytokine responsiveness of Mll1KO T cells is a consequence, rather than a cause, of impaired TOX and BTLA expression.

To understand how defects in activated Mll1KO T cells develop, I performed time-course analyses following T cell activation. Both WT and Mll1KO T cells downregulated TOX after activation; however, Mll1KO cells exhibited a more rapid and sustained reduction (Figure 5E). Differences were detectable as early as day 1 and persisted throughout the 3-day period. BTLA expression followed a different pattern, with initial upregulation followed by progressive downregulation (Figure 5F). Notably, the initial upregulation was not affected by MLL1 deficiency. However, the downregulation occurred earlier and more markedly in Mll1KO cells. These observations are consistent with BTLA being a downstream target of TOX ^72^. In addition to BTLA, TOX has been reported to regulate PD-1 (encoded by Pdcd1) expression. However, RNA-seq analysis indicated that *Pdcd1* transcription was not affected by MI-3454 treatment. Consistent with this, flow cytometry revealed similar PD-1 expression kinetics in WT and Mll1KO T cells, with sustained high expression in both groups (Figure 5G).

BTLA functions as a co-inhibitory receptor that restrains T cell activation, primarily through recruitment of the phosphatase SHP-1. Although SHP-1–independent effects have been reported, SHP-1 is required for sustained BTLA-mediated inhibition ^75^. In T cells, SHP-1 limits TCR and cytokine signaling by inhibiting PI3K, thereby reducing AKT activities ^76^. PI3K generates PIP3 from PIP2, creating a docking site for AKT activation by PDK1 and mTORC2 ^77^. Additionally, SHP-1 can attenuate cytokine signaling by dephosphorylating JAKs and STATs ^78^. Collectively, these findings establish TOX as a key downstream target of MLL1 and a central regulator of cytokine responsiveness in activated T cells.

### MLL1 regulates TOX independently of H3K4me3 and H4K16ac

To determine whether MLL1 regulates *Tox* transcription through its methyltransferase activity, I activated CD8⁺ T cells in the presence of WDR5 inhibitors (WDR5i or MM-401), which disrupt the MLL1–WDR5 interaction and block H3K4 methylation. As shown in Figure 6A, neither inhibitor altered *Tox* expression, indicating that MLL1 regulates *Tox* independently of its canonical methyltransferase activity. The *Tox* locus is regulated by both its promoter and multiple enhancer elements ^15,71,79^. To assess whether MLL1 influences H3K4me3 at this locus, I performed ChIP-PCR analysis at the transcription start site (+85) and previously identified enhancer regions (+22 kb, +50 kb, +90 kb, +110 kb, +122 kb, +133 kb, and +310 kb) ^47^. A gene desert region on chromosome 6 (Ch6-N) served as a negative control. H3K4me3 was highly enriched at the *Tox* promoter compared to enhancers and the control region; however, enrichment levels were comparable between Mll1KO and wild-type (WT) T cells (Figure 6B). These findings indicate that MLL1 does not regulate H3K4me3 at the *Tox* locus in activated T cells. *Tox* is expressed in hematopoietic stem cells and common lymphoid progenitors ^80^, downregulated during B cell development ^81^, and upregulated during T cell development ^73^. Consistent with this, I observed that *Tox* expression was ∼100-fold higher in thymocytes than in B cells (Figure 6C). However, ChIP-PCR analysis revealed that H3K4me3 enrichment at the *Tox* promoter was present in both cell types, with less than a threefold difference between them (Figure 6D). This disparity suggests that H3K4me3 does not directly instruct transcriptional output at the *Tox* locus as in many other gene loci ^50^. Instead, H3K4me3 may reflect prior transcriptional activity rather than serving as a primary regulatory determinant.

**Figure 6.**
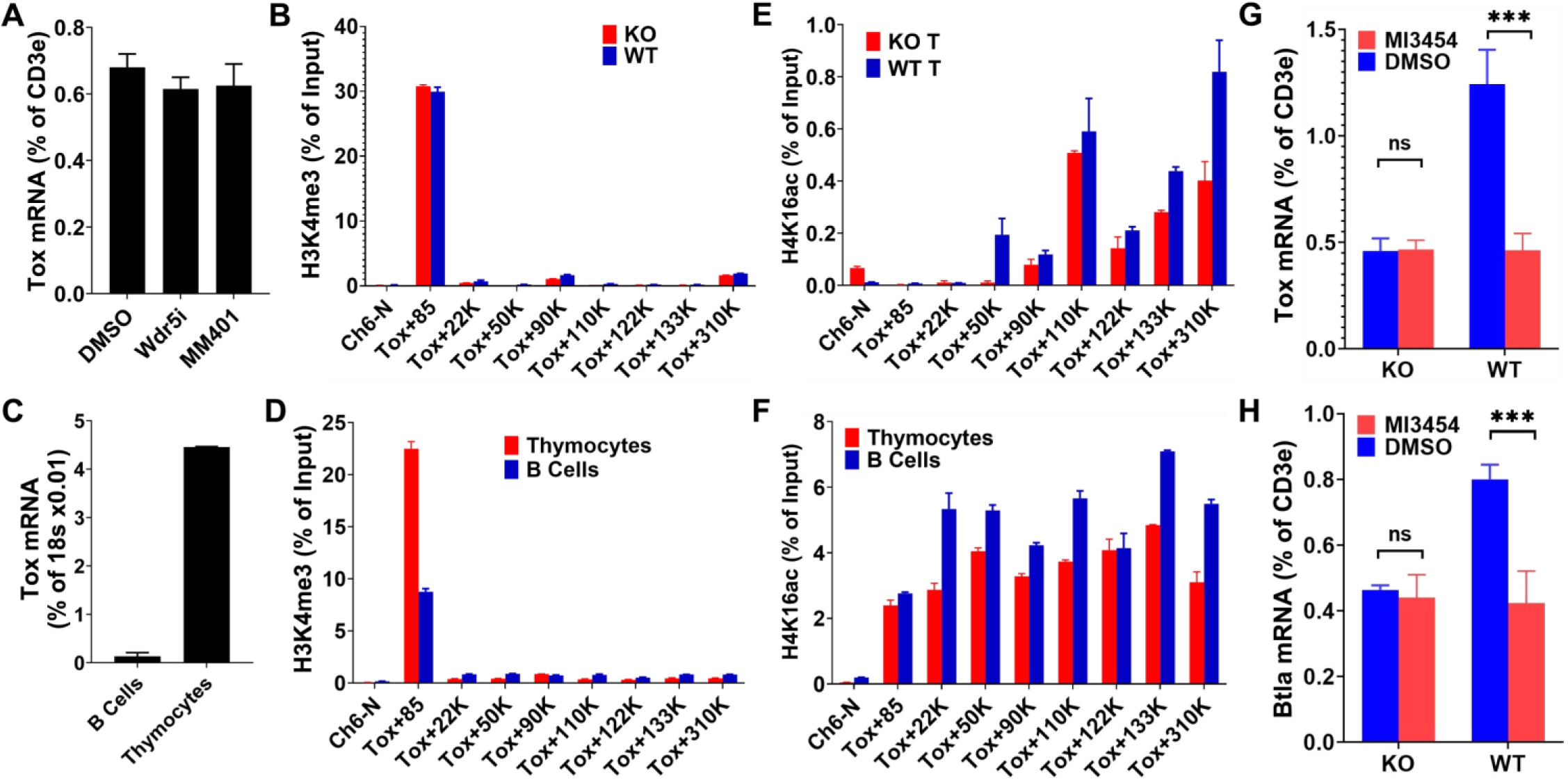
MLL1 regulates TOX independently of H3K4me3 and H4K16ac. (A) Wild-type (WT) T cells were activated in the presence of WDR5-IN-4, MM-401, or DMSO. After 4 days, cells were collected and analyzed for *Tox* expression by RT-PCR. Data are representative of two independent experiments. (B, E) T cells from Mll1KO mice and their WT littermates were activated in vitro. After 4 days, cells were collected and analyzed for H3K4me3 (B) and H4K16ac (E) enrichment at the *Tox* locus by ChIP-PCR. Data are representative of three independent experiments. (C, D, F) Thymocytes and B cells were isolated from WT mice and compared for *Tox* expression by RT-PCR (C), and for H3K4me3 (D) and H4K16ac (F) enrichment at the *Tox* locus by ChIP-PCR. Data are representative of two independent experiments. (G, H) T cells from Mll1KO mice and their WT littermates were activated in the presence of MI-3454 or DMSO. After 4 days, cells were collected and analyzed for *Tox* (G) and *Btla* (H) expression by RT-PCR. Data are representative of two independent experiments.

MLL1 has also been reported to regulate gene expression by recruiting the histone acetyltransferase MOF, leading to H4K16 acetylation (H4K16ac) at gene promoters ^28,29^. To assess the role of this pathway, I examined H4K16ac levels at the *Tox* locus. Contrary to prior reports, H4K16ac was not enriched at the *Tox* promoter and instead displayed a broad distribution across the locus (Figure 6E). Furthermore, H4K16ac levels were similar between Mll1KO and WT T cells. Next, I compared H4K16ac enrichment at the *Tox* locus in thymocytes and B cells. Despite the marked (∼100-fold) difference in *Tox* expression between these cell types, H4K16ac levels were comparable (Figure 6F). These findings suggest that H4K16ac does not play a major role in regulating *Tox* transcription in these contexts. Notably, while early studies reported an interaction between MLL1 and MOF ^29^, subsequent work has questioned this association ^82–84^. Although H4K16ac is enriched at the promoters in mouse embryonic stem cells, in most cell types, H4K16ac is broadly distributed outside promoter regions ^83^, consistent with my observations.

Finally, to determine whether MLL1 regulates *Tox* through interaction with MENIN, I activated WT and Mll1KO T cells in the presence of MI-3454 or DMSO and measured *Tox* and *Btla* expression by RT-PCR. In WT T cells, MI-3454 treatment reduced *Tox* (Figure 6G) and *Btla* (Figure 6H) expression. In contrast, MI-3454 had minimal effects in Mll1KO T cells, indicating that MLL1 and MENIN function cooperatively to sustain *Tox* transcription. Together, these results demonstrate that MLL1 maintains *Tox* (and *Btla*) transcription through interaction with MENIN, independently of both its methyltransferase activity and MOF-mediated H4K16 acetylation.

### MLL1 is required for T cells to induce graft-versus-host disease

Similar to Mll1KO T cells, BTLA-deficient T cells exhibit impaired ability to reconstitute lymphopenic hosts. In addition, BTLA-deficient T cells fail to induce graft-versus-host disease (GVHD) ^85–87^. Given that MLL1 is required to maintain BTLA expression in activated T cells, I hypothesized that MLL1 is also required for T cells to mediate GVHD. To test this, I employed a model of lethal GVHD ^88^. T cells from Mll1KO mice or their wild-type (WT) littermates were co-transferred with Rag1KO bone marrow cells into irradiated allogeneic B6D2F1 recipients (Figure 7A). As expected, all the mice receiving WT T cells succumbed to disease within one month (Figure 7B). In contrast, none of the mice receiving Mll1KO T cells developed lethal GVHD. These findings demonstrate that MLL1 is required for T cells to induce GVHD.

**Figure 7.**
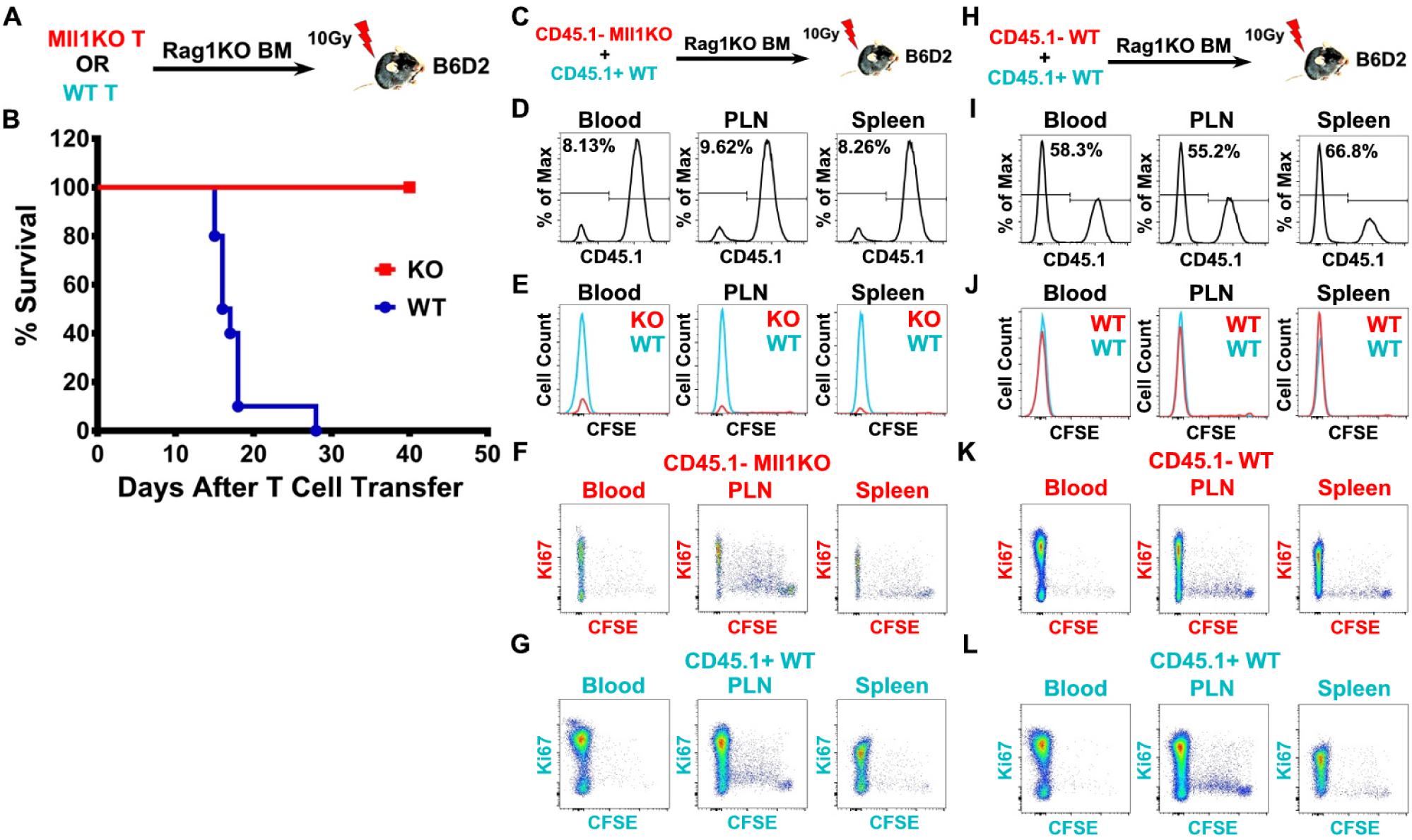
MLL1 is required for T cells to induce graft-versus-host disease. (A, B) CD8⁺ T cells from Mll1KO mice or their wild-type (WT) littermates were mixed with bone marrow cells from Rag1KO mice and transferred into lethally irradiated allogeneic B6D2F1 recipients. (A) Schematic of the experimental design. (B) Survival of recipient mice. Data are representative of two independent experiments. (C–G) Congenically marked CD8⁺ T cells from Mll1KO (CD45.1⁻ CD45.2⁺) and WT (CD45.1⁺ CD45.2⁻) mice were mixed at a 1:1 ratio, labeled with CFSE, and co-transferred with Rag1KO bone marrow cells into lethally irradiated allogeneic B6D2F1 recipients. (C) Schematic of the experimental design. (D–G) Ten days after transfer, donor T cells were analyzed by flow cytometry for the frequency of Mll1KO (CD45.1⁻) cells (D), CFSE dilution (E), and Ki-67 expression in Mll1 (F) and WT (G) donor T cells. Data are representative of three independent experiments. (H–L) Control experiments were performed using CD8⁺ T cells from WT (CD45.1⁻ CD45.2⁺) littermates of Mll1KO mice and congenically marked WT CD8⁺ T cells (CD45.1⁺ CD45.2⁻). Donor cells were mixed at a 1:1 ratio, labeled with CFSE, and co-transferred with Rag1KO bone marrow cells into lethally irradiated allogeneic B6D2F1 recipients. (H) Schematic of the experimental design. (I–L) Ten days after transfer, donor T cells were analyzed for the frequency of CD45.1⁻ cells (I), CFSE dilution (J), and Ki-67 expression in CD45.1⁻ WT (K) and CD45.1⁺ WT (L) donor T cells. Data are representative of two independent experiments.

To further investigate this defect, I co-transferred equal numbers of congenically marked CD8⁺ T cells from Mll1KO (CD45.1⁻) and WT (CD45.1⁺) mice into irradiated B6D2F1 recipients (Figure 7C). Previous studies have suggested that MLL1 is required for cell division due to its roles in regulating cell cycle–associated genes, S-phase checkpoint control, chromosomal alignment, and spindle assembly ^89–92^. To assess proliferation in vivo, CD8⁺ T cells were labeled with CFSE prior to transfer. Six days after transfer, large numbers of WT CD8⁺ T cells (CD45.1⁺) were detected in the blood and spleen of recipient mice (Figure 7D). By contrast, very few Mll1KO CD8 T cells (CD45.1-) were found in the recipients. Despite this, both WT and Mll1KO T cells underwent robust proliferation, as evidenced by complete CFSE dilution (Figure 7E) and sustained Ki-67 expression (Figure 7F, G). These results indicate that antigen-driven T cell proliferation can occur independently of MLL1, despite prior reports implicating MLL1 in cell cycle regulation.

To determine whether the loss of Mll1KO T cells was due to allogeneic rejection, I performed control co-transfer experiments using CD8 T cells from WT (CD45.1⁻) littermates of Mll1KO mice and congenically marked WT T cells (CD45.1⁺). No differences in expansion or persistence were observed between the two WT populations (Figure 7H–L), indicating that the loss of Mll1KO T cells is not attributable to rejection by the host. Together, these findings demonstrate that MLL1 is intrinsically required for the persistence and function of alloreactive T cells during GVHD. These data further support a model in which MLL1 sustains allogeneic T cell responses by restraining activation through a MLL1–TOX–BTLA regulatory axis.

### MLL1 limits the expansion of virtual memory T cells

Virtual memory (VM) T cells are antigen-inexperienced cells that arise from naïve T cells in response to cytokine signaling ^19,20^. Given that Mll1KO T cells exhibit enhanced cytokine responsiveness, I hypothesized that loss of MLL1 would promote expansion of the VM T cell compartment. Consistent with this idea, increased memory CD8⁺ T cell populations have been reported in both BTLA-deficient ^93^ and MENIN-deficient ^34^ mice, although the underlying mechanism and the relative contributions of VM versus true memory (TM) cells remained unclear. To investigate the role of MLL1 in memory T cell homeostasis, I compared CD8⁺ T cell populations in naïve Mll1KO mice and their WT littermates. In young (3-month-old) mice, the proportion of memory (CD44⁺) CD8⁺ T cells was approximately twofold higher in Mll1KO mice compared to WT controls (Figure 8A). Similar to observations in BTLA- and MENIN-deficient mice, the majority of these memory cells exhibited a central memory phenotype (CD62L⁺) (Figure 8B).

**Figure 8.**
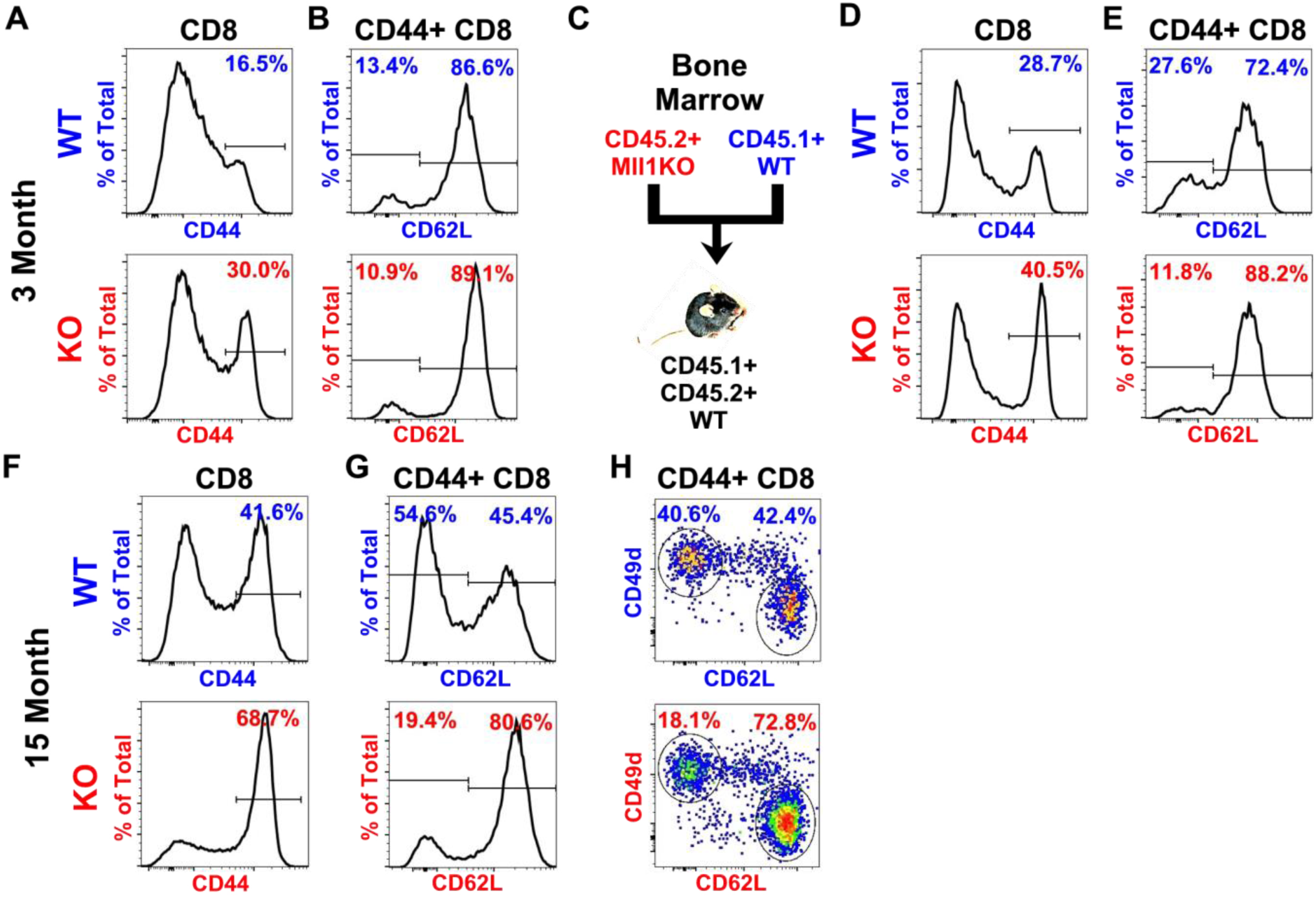
MLL1 limits the expansion of virtual memory T cells. (A, B) Splenocytes from young (3-month-old) Mll1KO mice and their wild-type (WT) littermates were analyzed by flow cytometry. (A) Gated CD8⁺ T cells showing the frequency of CD44⁺ (memory) cells. (B) Gated memory CD8⁺ T cells showing the frequency of CD62L⁺ (central memory) cells. Data are representative of five independent experiments. (C–E) Bone marrow cells from Mll1KO (CD45.1⁻ CD45.2⁺) and congenically marked WT (CD45.1⁺ CD45.2⁻) mice were mixed and transferred into lethally irradiated congenic WT (CD45.1⁺CD45.2⁺) recipients. (C) Schematic of the bone marrow chimera experiment. (D, E) Five months after reconstitution, splenocytes from recipient mice were analyzed by flow cytometry. (D) Gated CD8⁺ T cells showing the frequency of CD44⁺ (memory) cells. (E) Gated memory CD8⁺ T cells showing the frequency of CD62L⁺ (central memory) cells. Data are representative of three independent experiments. (F–H) Splenocytes from aged (15-month-old) Mll1KO mice and their WT littermates were analyzed by flow cytometry. (F) Gated CD8⁺ T cells showing the frequency of CD44⁺ (memory) cells. (G) Gated memory CD8⁺ T cells showing the frequency of CD62L⁺ (central memory) cells. (H) Gated memory CD8⁺ T cells showing the frequency of virtual memory (CD62L⁺ CD49d⁻) and true memory (CD62L⁻ CD49d⁺) subsets. Data are representative of two independent experiments.

To determine whether this expansion was cell-intrinsic, I generated mixed bone marrow chimeras using congenically marked Mll1KO (CD45.1⁻CD45.2⁺) and WT (CD45.1⁺CD45.2⁻) donor cells. Lethally irradiated CD45.1⁺CD45.2⁺ WT mice were used as recipients (Figure 8C). Five months after reconstitution, donor-derived T cells were analyzed by flow cytometry. Congenic markers enabled clear discrimination of residual host cells, Mll1KO, and WT donor populations. As shown in Figure 8D, Mll1KO CD8⁺ T cells exhibited a significantly higher proportion of memory cells compared to WT cells within the same host. The majority of these Mll1KO memory cells displayed a central memory phenotype (Figure 8E), and their frequency was higher than that observed in WT memory T cells. These findings indicate that the expansion of central memory T cells in Mll1KO mice is driven by cell-intrinsic mechanisms.

To assess the impact of aging, I analyzed aged (15-month-old) Mll1KO mice and WT littermates. As expected, the memory T cell compartment expanded with age (compare Figure 8A and 8F). Notably, the proportion of memory CD8⁺ T cells remained significantly higher in Mll1KO mice compared to WT controls. Moreover, the fraction of central memory (CD62L⁺) cells was consistently elevated in Mll1KO mice (Figure 8G). Previous studies have shown that central memory CD8⁺ T cells in naïve mice are predominantly VM cells ^21^. Unlike TM cells, which upregulate CD49d during antigen-driven differentiation, VM cells are characterized by low CD49d expression ^19^. To determine whether the expanded central memory population in Mll1KO mice corresponds to VM cells, I analyzed CD49d expression. In both Mll1KO and WT mice, central memory (CD62L⁺) CD8⁺ T cells expressed low levels of CD49d, whereas non-central memory (CD62L⁻) cells expressed high levels (Figure 8H). These results confirm that the expanded central memory population in Mll1KO mice consists predominantly of VM T cells. Together, these findings demonstrate that MLL1 restrains the expansion of virtual memory T cells, likely through maintenance of BTLA expression and control of cytokine responsiveness.

## Discussion

Immunological memory can persist for decades and provides durable protection against reinfection, yet the same durability can contribute to chronic inflammatory and autoimmune diseases. Understanding how memory T cells are maintained is therefore of both fundamental and clinical importance. The development and persistence of long-lived memory T cells depend on transcriptional programs that restrain terminal differentiation, among which TOX and TCF1 play central roles. In this study, I identify MLL1 as a critical regulator of this process and define a previously unrecognized pathway linking epigenetic regulation to cytokine signaling through a MLL1–TOX–BTLA axis.

A central and unexpected finding of this study is that MLL1 regulates *Tox*, *Tcf7*, and *Btla* independently of its canonical methyltransferase activity. MLL1 has been extensively studied as a histone methyltransferase responsible for H3K4 trimethylation ^94^. Most previously described defects in MLL1-deficient T cells have been attributed to loss of this activity ^95–97^. However, several lines of evidence indicate that the SET domain of MLL1 is dispensable for many of its biological functions ^98^, including stem cell maintenance ^28^. Consistent with these observations, my data show that MLL1 maintains the transcription of *Tox*, *Tcf7*, and *Btla* independently of its methyltransferase activity. Pharmacological inhibition of the MLL1–WDR5 interaction did not recapitulate the effects of MLL1 deficiency, and H3K4me3 levels at the *Tox* and *Tcf7* loci were not altered in MLL1-deficient T cells. Together, these findings indicate that, in activated T cells, MLL1 regulates key transcriptional programs through a noncanonical mechanism. MLL1 has also been proposed to regulate transcription through recruitment of MOF and H4K16 acetylation ^28,29^. However, recent studies have questioned the generality of this mechanism ^82–84^. Consistent with these reports, I found that H4K16ac was not enriched at the promoter of the *Tox* gene and did not differ between WT and MLL1-deficient T cells. Instead, my findings are consistent with a model in which MLL1 functions as a transcriptional cofactor, potentially through its ability to associate with MENIN and RNA polymerase II at actively transcribed loci ^25,26,99,100^. These data suggest that MLL1 maintains *Tox* transcription independently of both its methyltransferase activity and MOF-mediated H4K16 acetylation.

MLL1 has also been proposed to function as a mitotic bookmark that facilitates the re-establishment of transcription following cell division ^30^. My data are consistent with this model. The effects of MLL1 on *Tox, Tcf7* and *Btla* transcription are modest at the level of expression but have profound functional consequences. In activated T cells, where *Tox* and *Tcf7* are progressively downregulated, even small changes in transcriptional maintenance may critically influence cell fate decisions. More broadly, my findings suggest that the contribution of MLL1 to gene regulation is context dependent. Although MLL1 occupies a large fraction of actively transcribed promoters ^25,26^, its functional impact may be most apparent at genes expressed at low or dynamically regulated levels ^96,101^, such as *Tox* and *Tcf7* in activated T cells. In this setting, MLL1 appears to act as a stabilizer of transcriptional programs that are otherwise susceptible to cytokine-driven repression. Recent studies have shown that fully differentiated effector T cells can re-acquire memory-like properties through de-differentiation ^102,103^. The observation that H3K4me3 remains enriched at the *Tox* and *Tcf7* promoters even after transcriptional silencing raises the possibility that MLL1-mediated bookmarking contributes to this plasticity. It will be of particular interest to determine whether MLL1 is required for re-expression of these genes and for the de-differentiation of effector T cells.

My findings further provide insight into how cytokine signaling is integrated with transcriptional control of T cell differentiation. Cytokines such as IL-2, IL-4, and IL-12 promote effector differentiation in part through activation of AKT ^6,68–70^, which suppresses FOXO1 and downstream targets including *Tcf7* and *Sell*. I show that MLL1 limits this process by maintaining TOX and BTLA expression, thereby restraining cytokine responsiveness upstream of AKT. In this framework, BTLA act as key regulators of memory potential, while MLL1 functions to preserve their expression through TOX under conditions of activation. The role of BTLA in T cell biology has been difficult to reconcile, as BTLA deficiency enhances T cell-mediated inflammatory responses in some contexts ^31,104,105^ while impairing sustained T cell responses in others ^87,106^. My model provides a unifying explanation for the seemingly paradoxical roles of BTLA in T cell biology. While BTLA constrains effector responses, it also preserves memory potential by limiting cytokine-driven differentiation. Under conditions of strong or chronic stimulation, as in GVHD and colitis model ^87,106^, BTLA preserves a pool of T cells with memory potential by restraining cytokine-driven differentiation. Conversely, in settings where cytokine signals are limiting, as in EAE and allergen model ^31,104,105^, loss of BTLA may enhance effector responses. This finding has important implications for immunotherapy, as targeting co-inhibitory pathways may enhance effector responses at the cost of long-term memory. Finally, the persistent accumulation of large numbers of memory T cells in naïve BTLA-deficient mice has been difficult to reconcile with the fact that activated BTLA-deficient T cells are short-lived ^87,106^. My results suggest that the expanded memory population observed in naïve BTLA-deficient mice likely consists of virtual memory (VM) T cells, driven by enhanced cytokine responsiveness. Thus, co-inhibitory pathways may play a broader role in maintaining T cell homeostasis.

Although Mll1 is constitutively transcribed, MLL1 protein levels are dynamically regulated during the cell cycle ^107^, and its abundance in primary cells is relatively low ^108,109^. Due to the low abundance and dynamic regulation of endogenous MLL1 protein in primary T cells, I was unable to reliably detect differential MLL1 occupancy at the *Tox* promoter between WT and *Mll1*-deficient cells. Thus, I cannot formally exclude indirect contributions. However, several observations argue that MLL1 regulates *Tox* in a relatively direct manner, including the rapid transcriptional effects observed upon disruption of the MLL1–MENIN interaction and the concordant phenotypes between genetic and pharmacological perturbations. Notably, disruption of the MLL1–MENIN interaction has been explored in cancer, and my data suggest that such interventions may also modulate T cell differentiation and memory. Targeting this pathway could provide a means to either enhance effector responses or limit pathogenic memory T cells, depending on the context.

In summary, I identify a noncanonical function of MLL1 in maintaining *Tox* transcription through interaction with MENIN, thereby restraining cytokine signaling and preserving CD8 T cell memory. This work provides a conceptual advance by uncoupling MLL1 function from its widely assumed dependence on histone modification and identifying a mechanism by which modest transcriptional maintenance can have profound effects on T cell fate. Given the central importance of T cell memory in infection, autoimmunity, and immunotherapy, these findings may have broad implications for strategies aimed at modulating immune responses.

## Materials and Methods

### Mice

MLL1^flox/flox^ mice have been described previously ^33^. CD4-driven Cre transgenic mice STOCK Tg(Cd4-cre)1Cwi/BfluJ (Stock No: 017336) and Foxp3-driven Cre transgenic mice B6.129(Cg)-Foxp3^tm4^(YFP/icre)^Ayr^/J (Stock No: 016959) were purchased from the Jackson Laboratory (Bar Harbor, ME). Both strains were on the C57BL/6 (CD45.1⁻ CD45.2⁺) mouse background. T cell-specific MLL1-deficient (Mll1KO) mice were generated by crossing the MLL1^flox/flox^ with STOCK Tg(Cd4-cre)1Cwi/BfluJ mice. Treg cell-specific MLL1 deficient mice were generated by crossing the MLL1^flox/flox^ with B6.129(Cg)-Foxp3^tm4^(YFP/icre)^Ayr^/J mice. WT C57BL/6 congenic mice C57BL/6J (CD45.1⁻ CD45.2⁺) (Stock No: 000664) and B6.SJL-Ptprc^a^ Pepc^b^/BoyJ (CD45.1⁺ CD45.2⁻) (stock NO: 002014) were purchased from the Jackson Laboratory. C57BL/6 F1 congenic mice (CD45.1⁺CD45.2⁺) were produced by crossing male B6.SJL-Ptprc^a^ Pepc^b^/BoyJ (CD45.1⁺ CD45.2⁻) with female C57BL/6J (CD45.1⁻ CD45.2⁺) mice. Rag1 deficient mice B6.129S7-Rag1^tm1Mom^/J (Stock No: 002216) and B6D2F1/J (stock No: 100006) mice were purchased from the Jackson Laboratory. Mice were individually identified by ear tags and maintained under specific pathogen-free conditions and provided food and water ad libitum. The University of Michigan Committee on Use and Care of Animals (UCUCA) approved all animal studies.

### Reagents

Small-molecule inhibitors used included AKT inhibitor MK-2206 (selleckchem) used at 0.05uM, AKT inhibitor AKTi-1/2 (selleckchem) used at 0.5uM, Menin inhibitor MI-3454 ^37^ used at 0.25uM, Wdr5 inhibitor WDR5-IN-4 (Medchemexpress) used at 2.5uM, Wdr5 inhibitor MM-401 (invivochem) used at 25uM, Thymidine (for S-phase arrest) used at 2mM (sigma-aldrich), Nocodazole (for mitotic arrest) used at 0.5uM (selleckchem).

Antibodies used for ChIP-PCR included rabbit anti-H3K4me3 pAb from abcam ab8580 used at 2 µg per ChIP, rabbit mAb from CST 39751 (C42D8), use at 1:50 dilution (2ul), rabbit H3K27me3 mAb from abcam ab6002 used at 2ul per ChIP, rabbit H4K16ac mAb from abcam ab109463 used at 2ul per ChIP, Rabbit IgG from CST used at 2ul per ChIP, Anti-MLLc rabbit polyclonal from active motif (61295) used at 5-10ug (5-10ul) per ChIP, anti-MLLn rabbit mAb (D2M7U) from CST used at 1:50 dilution (2ul) per ChIP, anti-MLLc rabbit mAb (D6G8N) from CST used at 2ul per ChIP, anti-MLLn rabbit polyclonal from BETHYL (A300-086A) uses at 2-10ug (2-10ul) per ChIP.

Antibodies used for flow cytometry included anti-TCF1 (C63D9) from Cell Signaling Technology, anti-TCF1 (S33-966) from BD Pharmingen, phospho-S6 ribosomal Protein (pSer235/236) (D57.2) from Cell Signaling Technology, anti-CD19 (6D5), anti-CD62L (MEL-14), anti-CD44 (IM7), anti-CXCR3 (CXCR3-173), anti-TCRβ (H57-597), anti-CD49d (9C10), anti-CD122 (TM-β1), anti-CD5 (53-7.3), anit-CD6 (J90-462), anti-CD69 (H1.2F3), anti-CD127 (A7R34), anti-H2Kb (28-8-6), anti-Ki-67 (SolA15), anti-CD4 (GK1.5), anti-CD11b/Mac-1 (M1/70), anti-CD45 (30-F11), anti-CD45.1 (A20), anti-CD45.2 (104) and anti-CD8b (H35-17.2). Monoclonal antibodies were purchased from Biolegend, San Diego, CA or Serotec Raleigh, NC or eBioscience San Diego, CA or Cell Signal Technology and included eFluor 450-, APC-, APC-Cy7, AF700, PerCP-Cy5.5-, FITC-, PE-, PE-Cy7- and PE-Cy5-conjugated antibodies.

### T cell activation in vitro

Single cell suspensions were prepared from freshly harvested spleens and lymph nodes by mechanical dissociation. After washing with cold RPMI, cells were resuspended in culture media with anti-CD3 (1ug/ml), anti-CD28 (1ug/ml) and IL-2 (10ng/ml) at approximately 5 million cells per ml. In some cases, CD4 or CD8 T cells were removed before culture using anti-CD8 or anti-CD4 and magnetic beads respectively. For experiments with 3 or more days of culture, T cell purification was not needed because after 3 days of culture, all live cells were T cells. For flow cytometry, T cell purification was not needed. For other experiments with 2 or fewer days of culture, naïve CD8 T cells were isolated before culture using biotinylated anti-B220, anti-MHCII, anti-CD11b, anti-NK1.1, anti-TCRd and anti-CD4 antibodies and magnetic beads. One million T cells were stimulated with 20ul Dynabeads Mouse T-Activator CD3/CD28 (Thermo Fisher Scientific) and IL-2 (10 ng/ml) in 1 ml of complete medium. After 2 days, culture medium was replaced with IL-2 (10ng/ml) only.

### Lymphocyte reconstitution

Single cell suspensions were prepared from freshly harvested spleens and lymph nodes of Mll1KO (CD45.1⁻ CD45.2⁺) and congenic WT (CD45.1⁺ CD45.2⁻) mice by mechanical dissociation, mixed at a 1 to 1 ratio, activated in vitro, labeled with CFSE and intravenously transferred to T and B cell-deficient Rag1 deficient mice. Lymphocytes from the recipient mice were analyzed at various time points by flow cytometry.

### Generation of mixed bone marrow chimeras

To generate mixed bone marrow chimeras, C57BL/6 F1 congenic mice (CD45.1⁺CD45.2⁺) were irradiated with a single dose of 10 Gy and used as recipient mice. Bone marrow cells from congenically marked Mll1KO and WT donors were mixed at a predetermined ratio and transferred to recipients by intravenous injection within 2 hours of the irradiation treatment. After 5 months, recipients were analyzed.

### Modeling acute graft versus host disease (GVHD)

To model acute GVHD, B6D2F1/J mice were irradiated with a single dose of 10 Gy and used as recipient mice. Bone marrow cells from B6.129S7-Rag1^tm1Mom^/J (Rag1 deficient) mice were mixed with T cells isolated from congenically marked WT and/or Mll1KO mice and transferred to recipients by intravenous injection within 2 hours of the irradiation treatment. To monitor cell division, T cells were labeled with CFSE before transfer. In these recipients, donor cells can be identified by their lack of H2Kd, which is expressed by the host.

### Cytometry

Flow cytometric analyses were conducted as previously described ^110^. Briefly, single cell suspensions were prepared from freshly harvested spleens and lymph nodes by mechanical dissociation. For peripheral blood, samples were treated with lysis buffer and washed before antibody staining. Approximately 1 million cells were surface-stained with fluorochrome-labeled antibodies in 5% BSA-PBS buffer. Stained cells were washed with 1% BSA-PBS buffer, fixed and permeated with the Foxp3 Transcription Factor Fixation/Permeabilization Concentrate and Diluent solutions kit (ThermoFisher Scienticf). Fixed and permed cells were then stained with fluorochrome-labeled antibodies in 5% BSA-PBS buffer. After washing, cells were resuspended in 300ul of 1% BSA-PBS buffer, acquired using a BD LSR II flow cytometer, and data were analyzed with FLowJo software (Tree Star).

### Semi-quantitative RT-PCR (qPCR)

For qPCR analysis, RNeasy Plus Mini Kit (Qiagen) was used to isolate RNA from cells or tissues. After isolation, the concentration of the RNA was measured by nanodrop. Five hundred nanograms of RNA was reverse transcribed in a 20ul reaction using the iScript RT Supermix from Bio-rad. After reverse transcription, the products were diluted 10 times with DNase free water and used to run Real-time PCR with mouse 18s or Gapdh primer reagent from Applied Biosystems (Carlsbad, CA) in a 20ul reaction. The sample concentrations were adjusted so that all samples have the same cDNA concentration based on the results of the Real-time PCR. Samples were then used for real-time RCP analysis using pre-made primer reagent from Thermo Fisher Scientific. The results were analyzed using the comparative C_T_ method (ΔC_T_) methodology as outlined by Applied Biosystems. For comparing T cells, expression levels were normalized by CD3e. For other cells, levels were normalized by 18s.

### ChIP-PCR analysis

Single-use, methanol-free formaldehyde (1ml) were purchased from Thermo Fisher Scientific (28906), diluted to 16% formaldehyde (w/v) with PBS and used immediately. To crosslink DNA to proteins, cell pellets were resuspended in 16% formaldehyde (w/v) at less than 20 million cells per ml and incubated 10 minutes at room temperature with constant mixing. After 10 minutes, glycerin was added to samples at a 1:10 ratio and incubated 5 minutes at room temperature with constant mixing. Cells were then washed 2 times with 20ml cold PBS. After washing, cells were resuspended in 1 ml of cold PBS with protease inhibitor (Holt PIC from Thermo Fisher Scientific) at a concentration of 5ul per ml and transferred to a 1.7 ml sample tube. Cells were centrifuged, and cell pellets were then used to generate chromatin fragments by sonication. Sonication process began by cell lysis in sonication cell lysis buffer (Cell Signaling Technology, 96529) and then nuclear lysis in nuclear lysis buffer (Cell Signaling Technology, 28778). Sonication was conducted using a Branson probe sonicator with samples emersed in ice water. To completely break the nuclei, samples were sonicated in 0.5 ml of the nuclear lysis buffer using a 15% amplitude, 1 second on, 1 second off cycle for a total of 30 second on time. To fragment the chromatin, lysed samples were sonicated in 0.5 ml of the nuclear lysis buffer using a 10% amplitude, 1 second on, 1 second off cycle for a total of 2 minutes on time. Sheared chromatin was collected by centrifugation. Twenty microliter of the samples were taken to determine the sizes of the sheared chromatin and the DNA concentration of the samples. The remaining 480ul of the samples were used for immunoprecipitation with antibodies against targets of interest. Samples were used for immunoprecipitation only when the size of most of the sheared chromatin was between 500 to 1000 base pairs. Chromatin immunoprecipitation was conducted using protein A/G magnetic beads from Thermo Fisher. After chromatin immunoprecipitation, DNA was purified and subjected to qPCR analysis. For H3K4me3 immunoprecipitation, 2 different antibodies were used (anti-H3K4me3 pAb from abcam ab8580 and rabbit mAb from CST 39751 (C42D8)). The promotor of the Actb gene and a gene desert region on chromosome 6 (Ch6-N) were used as positive and negative control respectively. The two antibodies yielded similar results. For MLL1 immunoprecipitation, 5 different antibodies were used (Anti-MLLc rabbit polyclonal from active motif (61295), anti-MLLn rabbit mAb (D2M7U) from CST, anti-MLLc rabbit mAb (D6G8N) from CST, anti-MLLn rabbit polyclonal from BETHYL (A300-086A)).

## Notes

### Competing Interest Statement

The authors have declared no competing interest.

